# Interaction of osteoprotegerin with fibulin-1 is essential for extracellular matrix deposition in fibrotic lung tissue

**DOI:** 10.1101/2024.12.05.626371

**Authors:** Y Liu, H. Habibie, Theo Borghuis, Gang Liu, Philip M Hansbro, B.N. Melgert, J.K. Burgess

## Abstract

Idiopathic pulmonary fibrosis (IPF) is a progressive, fatal lung disease with an unclear cause and no cure. Osteoprotegerin (OPG), a well-known regulator of bone extracellular matrix (ECM), is increased in pulmonary fibrosis and this is associated with lower lung function and IPF progression. Fibulin-1, a key ECM protein, is increased in lung fibroblasts and lung tissues from patients with IPF and high levels are predictive of a poor prognosis for these patients. We found a positive correlation between levels of OPG and fibulin-1 in serum from patients with IPF, but it is unknown if these proteins interact. This study aimed at investigating interactions between OPG and fibulin-1 in lung fibrosis, hypothesizing that OPG may form a complex with fibulin-1 thereby possibly influencing lung fibrosis.

We investigated intracellular interactions between OPG and fibulin-1 in human lung fibroblasts, showing neither OPG nor fibulin-1 expression were dependent on each other. However, OPG protein deposition was significantly decreased in lung tissues of fibulin-1 knockout mice (fibulin-1^−/−^)compared to controls suggesting extracellular interactions. Subsequently, we detected colocalization of both proteins in both IPF and control lung tissues using immunofluorescence, with both of them also colocalizing with latent TGFβ binding protein 1 (LTBP1). Proximity ligation analyses confirmed close proximity of OPG to fibulin-1 and fibulin-1 to LTBP1, particularly in the interstitial regions of both control and IPF lung tissue, but not OPG and LTBP1.

In conclusion, we found that OPG, fibulin-1 and LTBP1 seem to form a complex, with fibulin-1 playing a bridging role between OPG and LTBP1, in the extracellular environment. This complex may play a regulatory role in deposition of extracellular matrix in lung tissue.

## Introduction

Idiopathic pulmonary fibrosis (IPF) is a chronic, progressive disease characterized by an increase in activated fibroblasts that secrete excessive amounts of extracellular matrix (ECM) resulting in an abnormal lung tissue structure^1^. Exaggerated deposition of ECM components, such as collagens and fibronectin, in the lung interstitium impede gas exchange leading to breathlessness and loss of lung function^2^. Patients with IPF are typically over 60 years and have a median survival of 3-5 years after diagnosis^3^. Pirfenidone and nintedanib are two treatments currently approved for slowing the progression of IPF, however, neither of them reverse or cure the disease^4,5^. Therefore, novel clinical management approaches are required based on a thorough understanding of the pathological processes driving development of fibrosis.

Osteoprotegerin (OPG) is a protein that is a novel player in the pathogenesis of fibrotic disorders. It is a soluble decoy receptor for the cytokines receptor activator for nuclear factor κB ligand (RANKL) and TNF-related apoptosis-inducing ligand (TRAIL) and is best known as a protector of bone ECM degradation^6^. In bone, OPG binding to RANKL prevents osteoclast activation and thereby degradation of bone tissue. OPG expression is induced by a variety of cytokines, hormones and growth factors, including transforming growth factor beta 1 (TGFβ1), the master regulator of fibrosis ^7,8,9^.

We and others have recently shown that OPG is not solely responsible for regulating bone matrix, it also plays a critical role in the development of fibrotic disease in lung, liver and heart^10,11,12,13^. We recently showed OPG was expressed at higher levels in lung tissue and lung fibroblasts of patients with IPF as compared to controls and was induced by TGFβ treatment of precision-cut lung slices ^11,14^. Moreover, OPG production was associated with lower lung function and accelerated IPF progression^14,15^. Nevertheless, how OPG contributes to progression of pulmonary fibrosis is still an open question.

Interestingly, in our recent data we found a positive correlation between serum levels of OPG and fibulin-1 in patients with IPF. Fibulin-1 is a glycoprotein secreted by lung fibroblasts^16^, which binds to other ECM proteins, such as latent TGFβ binding protein 1 (LTBP1)^17^, fibronectin, collagen I and laminin, and promotes ECM stability^16,18^. Fibulin-1 expression is higher in fibroblasts and lung tissues from patients with IPF compared to controls and high levels are predictive of a poor prognosis for these patients^19^. Fibuin-1 comprises four polypeptides in humans (fibulin1a, -b, -c, -d) identifiable by their different carboxy terminal regions and only fibulin-1c and fibulin-1d have been identified in mice^20^. Among these variants, only fibulin-1c has been associated with respiratory disease. We previously found that in chronic obstructive pulmonary disease it plays a crucial role in collagen production via interactions with its binding proteins^21^. With the association between OPG and fibulin-1 in serum of patients with IPF, we hypothesized that OPG may be interacting with fibulin-1 in contributing to IPF pathogenesis. Therefore, in this study we aimed to investigate if OPG and fibulin-1 and combinations thereof colocalize and functionally influence production of ECM and development of pulmonary fibrosis.

## Materials and Methods

### Ethics statements

Human lung tissue for this study was obtained with consent from lungs of patients with IPF undergoing transplantation for their disease or from control patients undergoing lung resection surgery for suspected cancer. The study was conducted in accordance to the Research Code of the University Medical Center Groningen (UMCG), as stated on https://umcgresearch.org/w/research-code-umcg as well as national ethical and professional guidelines Code of Conduct for Health Research (https://www.coreon.org/wp-content/uploads/2023/06/Code-of-Conduct-for-HealthResearch-2022.pdf). The study was not subject to Medical Research Human Subjects Act in the Netherlands, as confirmed by a statement of the Medical Ethical Committee of the University Medical Center Groningen and therefore exempt from consent according to national laws (Dutch laws: Medical Treatment Agreement Act (WGBO) art 458 / GDPR art 9/ UAVG art 24). All donor material and clinical information were deidentified prior to experimental procedures, blinding any identifiable information to the investigators.

### Patient details

**Table 1:**
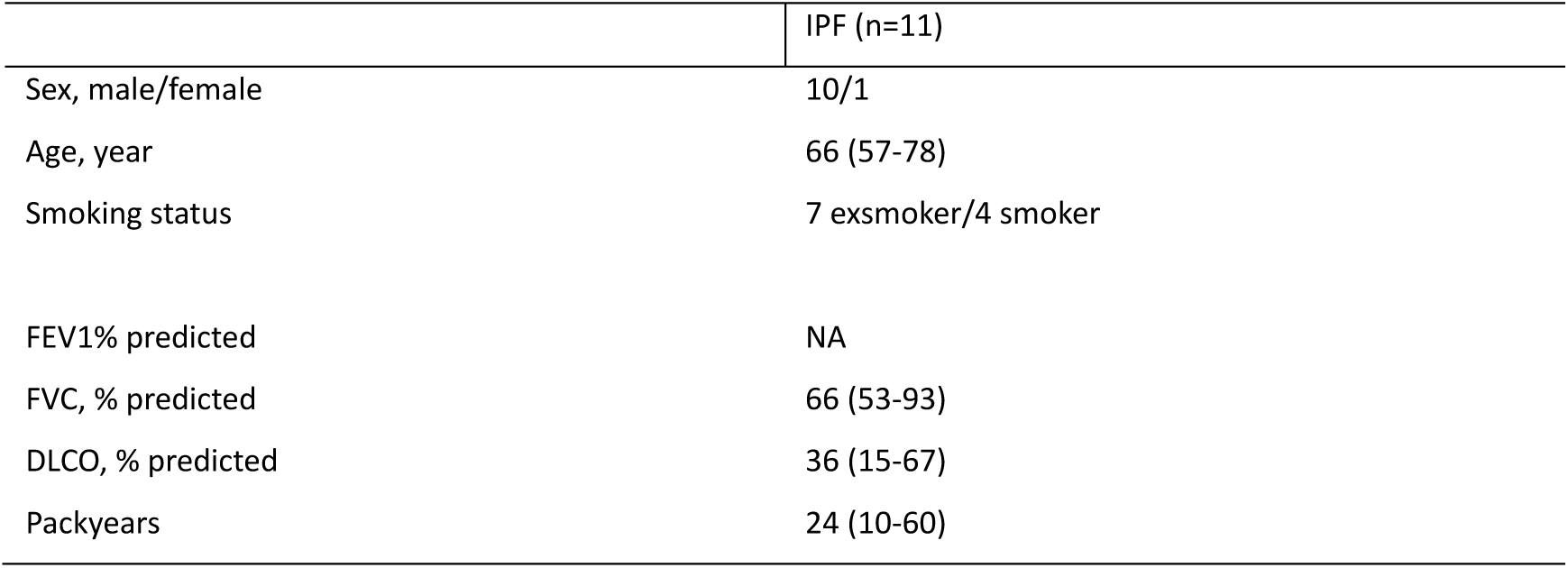
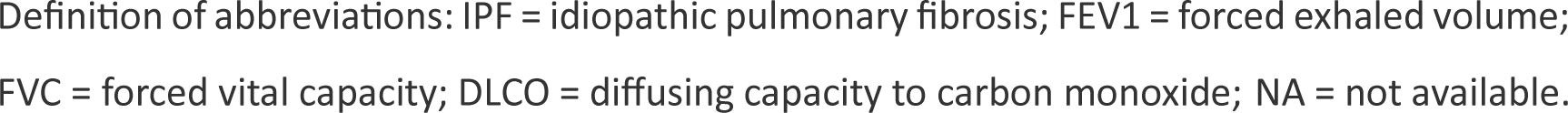
Patient characteristics of donors of serum used for OPG and fibulin1 measurements.

|  | IPF (n=11) |
| --- | --- |
| Sex, male/female | 10/1 |
| Age, year | 66 (57-78) |
| Smoking status | 7 exsmoker/4 smoker |
| FEV1% predicted | NA |
| FVC, % predicted | 66 (53-93) |
| DLCO, % predicted | 36 (15-67) |
| Packyears | 24 (10-60) |

Definition of abbreviations: IPF = idiopathic pulmonary fibrosis; FEV1 = forced exhaled volume; FVC = forced vital capacity; DLCO = diffusing capacity to carbon monoxide; NA = not available.

**Table 2:** Patient characteristics of donors of lung tissue used for immunofluorescent staining.

|  | Age (median) | Sex (male/female) | Number |
| --- | --- | --- | --- |
| Control | 55 (41-67) | 4/6 | 10 |
| IPF | 58 (39-67) | 5/5 | 10 |

Definition of abbreviations: IPF = idiopathic pulmonary fibrosis

**Table 3:** Patient characteristics of donors of lung tissue used for the proximity ligation assay.

|  | Age (median) | Sex (male/female) | Number |
| --- | --- | --- | --- |
| Control | 53 (45-72) | 3/2 | 5 |
| IPF | 60 (52-67) | 3/3 | 5 |

Definition of abbreviations: IPF = idiopathic pulmonary fibrosis

### Human lung fibroblasts isolation and culture

Primary human lung fibroblasts were obtained from lung tissue of ex-smoking control patients and patients with IPF as described previously^22^. The cells were seeded at a density of 15000 cells/cm^2^ in 6-well plate, cultured for 24 hours in Dulbecco’s modified Eagle medium-low glucose (DMEM, 31885-023, Thermo Fisher Scientific (Burlington, USA)) containing 10% fetal bovine serum (FBS) (Bovogen; Sigma–Aldrich, Zwijndrecht, the Netherlands), 1% GlutaMAX (Gibco), 100 U/mL penicillin, and 100 μg/mL streptomycin (Gibco, Breda, the Netherlands) (37°C, 5% CO2). After 24 hours, the medium was removed, and cells were ready for transfection with small interfering RNAs (siRNAs).

### Transfection with small interfering RNAs (siRNAs)

siRNAs inhibiting expression of OPG (TNFRSF11B targeting Silencer Select siRNA, 4390824, assayID s9858; Negative siRNA control, 4390843) were bought from Thermo Fisher Scientific (Burlington, USA) and those targeting fibulin-1 (Fibulin 1c siRNA, custom siRNA) were bought from Dharmacon (Baltimore, USA). siRNAs were dissolved using nuclease-free water to a concentration of 100uM. Targeting siRNAs were diluted with Opti-MEM™ I Reduced Serum Medium (31985062, Thermo Fisher Scientific) from 100uM to 20uM, then diluted separately from 20 uM to 2uM. siRNAs were mixed with Lipofectamine™ RNAiMAX Transfection Reagent (13778100, Thermo Fisher Scientific) gently to 5nM siRNA to become siRNA complexes, and equilibrated at room temperature for 10 min. After removing medium as mentioned above, cells were transfected with siRNA complexes 400 µL per well for 6 hours while a control condition was incubated in pure Opti-MEM medium with transfection reagent for same length of time at 37°C in an incubator. After that, the transfection medium was removed, and cells were treated with or without 1ng/mL TGFβ1 (made in Opti-MEM medium) for 24 hours. After 24 hours, TGFβ1-containing medium was replaced with fresh DMEM containing 10% FBS for 48 hours at 37°C in an incubator. After incubation, culture media were collected for OPG ELISA, cells were rinsed with cold PBS twice, cell lysates were isolated using TRIzol (1596026, Invitrogen) and collected for mRNA analysis. Radioimmunoprecipitation assay buffer (RIPA buffer) (Thermo Fisher Scientific) was used to collect cell lysates for Western blot analysis.

### mRNA isolation and quantification

After isolation of RNA, total RNA concentrations were measured using a NanoDrop (Thermo Fisher Scientific). cDNA was synthesized using TaqMan Gene Expression assays (Thermo Fisher Scientific) according to the manufacturer^’^s protocol: 5 min at 65°C, 5 min at 25°C, 60 min at 42°C, 5 min at 70°C. Quantitative polymerase chain reaction (qPCR) was performed in triplicate with the synthesized cDNA to quantify transcription of hypoxanthine-guanine phosphoribosyltransferase 1 (HPRT1), OPG, fibulin-1c, collagen-1a1 (Col1a1), fibronectin (Fn), fibulin-1d (fbln-1d), and Tgfb1 using a ViiA™ 7 real-time PCR system (Life Technologies). qPCR analysis consisted of 45 cycles of 10 minutes at 95°C, 15 seconds at 95°C, 25 seconds at 60°C (repeated for 40 times) followed by a dissociation stage of 95°C for 15 seconds, 60°C for 15 seconds, and 95°C for 15 seconds. Expression was quantified using QuantStudio™ Real-Time PCR Software (Thermo Fisher Scientific) and ΔCt values were normalized to the housekeeping gene HPRT1 and relative gene expression was calculated as 2^−ΔΔCt^.

### Western Blot

Human and mouse OPG and fibulin-1 expression was assessed using Western blot. Samples were diluted with 4x Laemmli protein sample buffer (BIO-RAD, California, USA) for SDS page and were denatured at 100 °C for 10 minutes. 20 μL samples were loaded onto 8% polyacrylamide gels and run for 30 min at 60 V first and then run at 120 V for 85 min. Proteins were transferred to nitrocellulose blot membranes using Trans-Blot Turbo Transfer System (Bio-Rad) 1.3A, 25V, for 30 min. To evaluate the transfer efficiency, membranes were stained using Ponceau S (Thermo Fisher Scientific) for visualization of bands. Ponceau S was removed afterwards by washing three times for a total of 5 minutes with 1X Tris-Buffered Saline, 0.1% Tween® 20 detergent (TBST). Aspecific binding was blocked with 5 % milk in TBST solution for 45 min. The membrane was then incubated overnight at 4 °C with a primary antibody against either OPG (1:1000 in 1% milk/TBST, ab183910, Abcam), OPG (1:2000, SCZ-sc-390518, Santa Cruz), fibulin-1 (1:750 in 1% ELK/TBST, ab211536, Abcam), or loading control β-actin (1:5000 in 1% milk/TBST, sc-47778, SantaCruz). The next day, membranes were washed with TBST three times for a total of 15 minutes. Subsequently, the membranes were incubated with secondary antibodies, goat anti-rabbit immunoglobulin-HRP (1:5000 in 1% milk/TBST, Dako P0488), or rabbit anti-mouse immunoglobulin-HRP, (1:5000 in 1% milk/TBST, Dako P0260) for 45 minutes. Thereafter, membranes with OPG and B-actin antibodies were washed with TBST three times for a total of 45 minutes. The membrane with fibulin-1 antibodies was washed with TBST three times for total 15 minutes and incubated with a tertiary antibody goat anti-rabbit immunoglobulin-HRP (1:5000 in 1% milk/TBST, Dako P0260) for 45 minutes and subsequently washed with TBST three times for a total of 45 min. For detection, all membranes were incubated in SuperSingal West Pico PLUS chemiluinescent substrate for 5 min (Thermo scientific, 34580). Membranes were placed and signals were captured in a ChemiDoc MP imaging system (Bio-Rad).

### Enzyme-linked immune sorbent assay (ELISA)

OPG expression in serum from patients with IPG, and in cell media were assessed using an human OPG ELISA kit (cat#DY805, R&D Systems Minneapolis) or a mouse OPG ELISA kit (cat#DY459, R&D Systems) according to the instructions provided by the manufacturer. Samples were diluted 1:100 (OPG) in reagent diluent.

### Mice and bleomycin-induced fibrosis

Wild type (WT) C57BL/6J (female, 6-8 weeks old) mice were purchased from Australian BioResources and *Fbln1c^−/−^* mice (6-8-weeks) were generated as previously described^21^.The mice were maintained in specific pathogen–free conditions and were bred in the Animal Facility at the Centenary Institute, Sydney. The mice received one dose of bleomycin sulfate (0.05 U/mouse, MP Biomedical) intranasally to induce the experimental model of pulmonary fibrosis as described previously^17,23^. Control mice received an equal volume of sterile PBS. Lung tissue was collected 28 days after bleomycin administration.

### Mouse lung fibroblasts isolation and culture

Primary lung fibroblasts were isolated from both WT and *Fbln1c^−/−^*mice as previously described. The fibroblasts (1×10^5^ cells) were seeded in 6 well plates with 10% FCS DMEM media for 24 hours. The media was then changed to low serum medium (0.1% BSA) for another 24 hours. Some cells then treated with 5 ng/mL TGFβ1 (cat#7666-MB, R&D Systems) and incubate for 24 hours. The cells were then used for western blot.

### Immunohistochemistry

Four-μm sections of paraffin-embedded lung tissues from mice were deparaffinized and antigens were retrieved through incubating in 10 mM citrate buffer at pH 6.0 at 100℃ for 15 min. Endogenous peroxidases were blocked by incubating in 0.3% H2O2 (Merck KGaA) at room temperature for 30 min. After washing three times in phosphate-buffered saline (PBS), sections were blocked with 4% BSA at room temperature for 30 min. Subsequently, sections were incubated with an OPG antibody (1:300, ab183910, Abcam) overnight at 4°C. Following washing 3 times, slides were incubated with goat anti-rabbit immunoglobulin-HRP (1:100, Dako P0488) for 45 min. After washing three times in PBS, slides were washed in demineralized water once. Vector®NovaRED® (Vector Laboratories, California, United States) substrate was then used to visualize the staining. After washing sections in demi water three times, sections were counterstained with hematoxylin, mounted and scanned using the 40x objective of a Hamamatsu NanoZoomer digital slides scanner (Hamamatsu Photonic K.K)

### Immunofluorescence

Four-μm sections of paraffin-embedded lung tissues from 10 patients with IPF and 10 control patients with normal lung function (see table 2 for patient characteristics) were deparaffinized and antigens were retrieved through incubating in 10 mM citrate buffer at pH 6.0 at 100℃ for 15 min. Endogenous peroxidases were blocked by incubating in 0.3% hydrogen peroxide (H2O2) (Merck KGaA, Darmstadt, Germany) solution at room temperature for 30 min. After washing three times in 1X Tris-Buffered Saline, 0.1% Tween® 20 Detergent (TBST), tissue sections were blocked with 4% bovine serum albumin (BSA) in TBST at room temperature for 30 min. Those sections were incubated with a primary antibody. Antibodies against OPG, fibulin-1 and LTBP1 were bought from Abcam, Cambridge, UK. Firstly, the fibulin-1 antibody was diluted in 1%BSA/TBST for fibulin-1 (1:100, ab211536, Abcam) incubated overnight at 4 ^°^C. Following washing, slides were treated with secondary antibody for 45 min. Rabbit anti-mouse immunoglobulin-HRP and goat anti-rabbit immunoglobulin-HRP were bought from Dako, Amsterdam, NL. Sections were incubated with rabbit anti-mouse immunoglobulin-HRP (1:100, P0260). The slides were washed three times in TBST, and subsequently incubated with tertiary goat anti-rabbit immunoglobulin-HRP (1:100, P0488) for 45 min. After a third wash step, slides were incubated with Opal fluorophore reagents (Akoya Biosciences, Marlborough, MA, US). Sections with the fibulin-1 antibody were incubated with Opal 650 reagent (1:200, FP1496001KT, Akoya Biosciences) diluted in 100mM borate buffer pH 8.5 containing 0.1% Tween-20, 0.003% H_2_O_2_ and for 15 min. Bound antibodies were subsequently removed by incubating slides in 10mM citrate buffer pH 6.0 for 15 min at 100 °C and the same steps as mentioned above were applied for blocking. After those, the sections were incubated with a primary antibody diluted in 1%BSA/TBST against OPG (1:300, ab183910, Abcam) overnight at 4 ^°^C. After washing, sections were incubated with secondary goat anti-rabbit immunoglobulin-HRP (1:100, Dako P0488,) for 45 min at room temperature. After a third wash step, slides were incubated with Opal 570 reagent (1:200, FP1488001KT, Akoya Biosciences) for 15 min. Again, bound antibodies were removed and blocking was done as described above. The sections were then incubated overnight at 4 °C with a primary antibody against LTBP1 diluted in 1%BSA/TBST (1:100, ab78294, Abcam,). Following washing, sections were incubated with secondary goat anti-rabbit immunoglobulin-HRP (1:200, Dako P0488) for 45 min at room temperature After washing, the slides were incubated with Opal 520 reagent (1:200, FP1488001KT, Akoya Biosciences) for 15 min. After that, sections were incubated with 4′,6-diamidino-2-phenylindole (DAPI) (Roche Diagnostics GmbH, Mannheim, Germany) for 10 minutes to visualize cell nuclei. Fluorescence signals were captured using the 20x objective of Olympus VS200 Slide Scanner. LTBP1, OPG and fibulin-1 signals were obtained at wavelengths 488nm, 594nm, and 633nm respectively.

### Proximity Ligation Assay

To show close colocalization of fibulin1, OPG and LTBP1 we used a NaveniBright HRP kit from Navinci (NB/MR.HRP.100, Uppsala, Sweden), which can visualize close proximity of these proteins. Four-um sections of lung tissue embedded in paraffin from 5 different control patients and patients with IPF (see table 1 for patient characteristics) were used. For each pair of proteins (OPG + fibulin-1, fibulin-1 + LTBP1, and OPG + LTBP1), slides were deparaffinized and rehydrated, followed by antigen retrieval with 10mM citrate buffer. Endogenous peroxidase activity was blocked by incubating the slides for 30 min in PBS containing 0.3% hydrogen peroxide at room temperature followed by overnight incubation at 4℃ with the primary antibodies of each combination. Fibulin-1 (monoclonal mouse, anti-human, sc-25281, 1:200, SantaCruz, California, USA) + OPG (polyclonal rabbit, anti-human, ab183910, 1:300, Abcam), fibulin-1 (sc-25281, 1:200) + LTBP1 (monoclonal mouse, anti-human, sc-271140, 1:75, SantaCruz) and LTBP1 (monoclonal mouse, anti-human, sc-271140, 1:75, SantaCruz) + OPG (ab183910, 1:300) were diluted in Tris-buffered saline + 2% BSA. The sections were then washed and incubated with secondary antibodies (PLA probe PLUS and MINUS, conjugated with oligonucleotides). This was followed by a 30 min period for enzymatic ligation at 37°C, which can only happen when the PLA probes PLUS and MINUS are physically located within 40 nm of each other^24^. Next, amplification was performed at 37°C for 60 min after which the slides were incubated in NaviBright HRP reagent for 30 min at room temperature to visualize the close proximity of the proteins, followed by staining nuclei with hematoxylin. As a negative control, we omitted either one of the two antibodies. Tissue sections were scanned with the 40x objective of a Hamamatsu NanoZoomer 2.0HT digital slide scanner (Hamamatsu Photonic K.K, Japan). Following scanning, digital images were viewed using Aperio ImageScope V 12.4.6 (Leica Biosystem, Nussloch, Germany). Figure 1 illustrates the process of proximity ligation assay staining.

**Figure 1:**
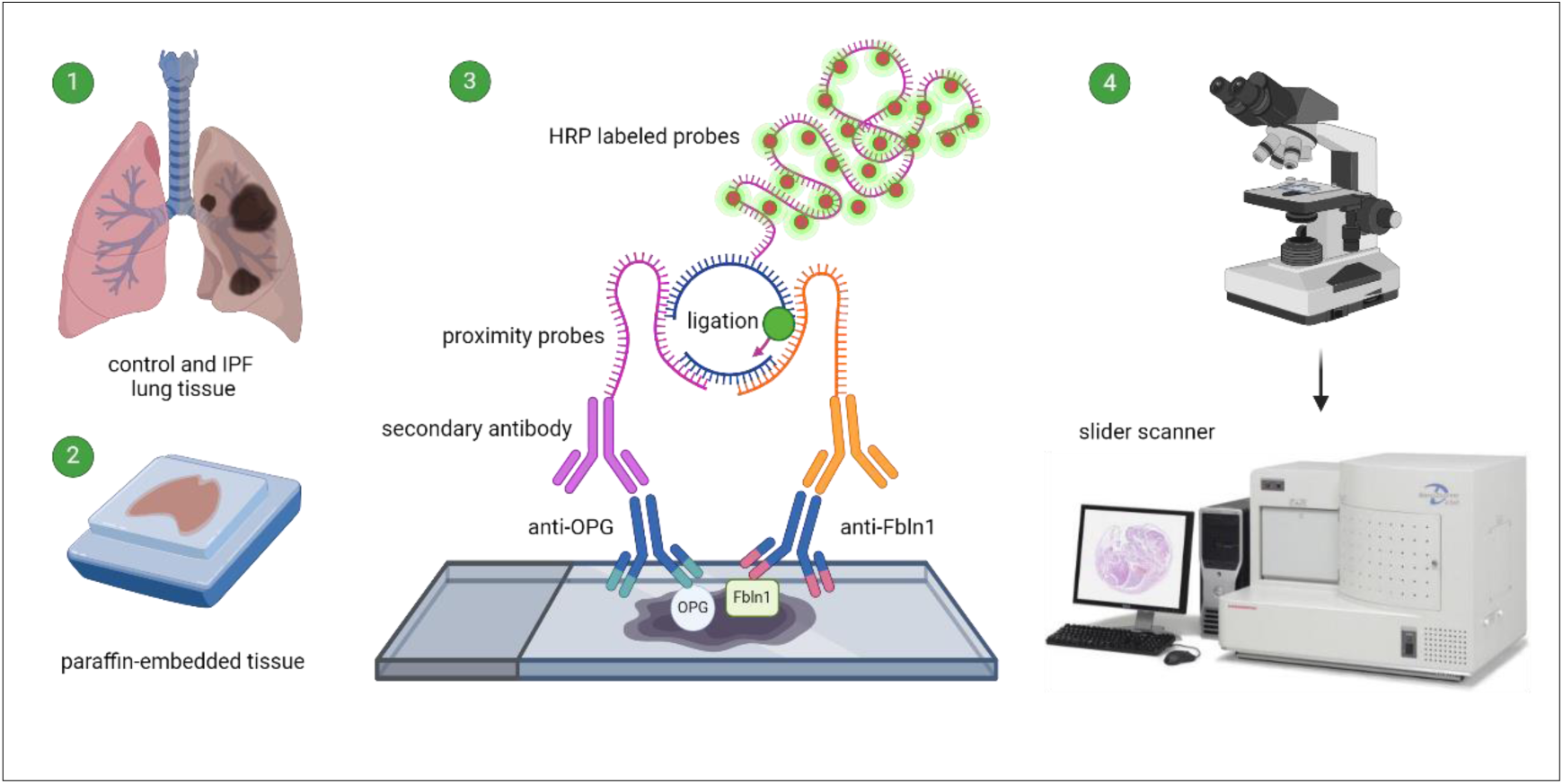
Tissue preparation, proximity ligation assay, and image analysis. Lung tissue from both controls and patients with IPF were embedded in paraffin and cut in sections. The sections were then stained for visualization of proximity of OPG and fibulin-1, fibulin-1 and LTBP1, as well as OPG and LTBP1. After the staining, positive signals were visualized with a brightfield microscope. The images were captured using a Hamamatsu NanoZoomer 2.0HT at a magnification of 40x.

### Imaging analysis

Images were extracted using Aperio ImageScope V 12.4.6 (Leica Biosystem) and staining artefacts (folded tissues and hairs) were removed using Adobe Photoshop 2024 (Adobe Inc. California, United States). The subsequent TIF images were then opened in FIJI^25^ (LOCI, University of Wisconsin) and color deconvolution was applied splitting the staining into different channels that represented different staining colors. We then selected 5 images per staining randomly to set up the thresholds for measurement of the amount of each color present in the images and used 8-bit images to represent the total tissue area. All thresholds were used in a macro calculating tissue area per channel. After running all images, signals of pixels were categorized based on levels of intensity values in R studio (Boston, MA, USA) and sorted for the percentage of pixels of positive stained area per image. The calculation of the percentage positively stained area was performed using equation as shown below.

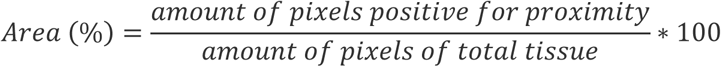

### Statistical analyses

Data sets with n ≤8 were considered nonnormally distributed and differences between multiple groups were therefore assessed using a nonparametric Friedman test for paired and Kruskal-Wallis test for unpaired data. A two-way analysis of variance (ANOVA) was used to identify the impact of bleomycin and fibulin-1 individually and to identify when these two conditions were interacting. A two-way ANOVA analysis allows to test whether exposure of bleomycin causes significant change compared to saline, whether the absence of fibulin-1 causes a significant change compared to the presence of fibulin-1, and whether these two conditions have significant interaction effect, meaning is the combination of the two conditions more or less than a simple condition. In case of significant interaction, the outcomes for the individual conditions cannot be trusted anymore and Fisher Least Significant Difference post-hoc test was utilized to compare individual groups to explain the interaction. P < 0.0.5 was considered significant. Correlations were assessed using a nonparametric (Spearman) correlation coefficient (r).

## Results

### Serum OPG and fibulin-1 levels correlate positively in patients with IPF

Previously we have shown that both OPG and fibulin-1 levels in serum independently correlate with disease severity in patients with IPF^14,19^. To investigate if these two proteins could be interacting in some way, we measured fibulin-1 serum levels in the same cohort in which we had previously assessed OPG levels^26^ and we found that OPG levels positively correlated with fibulin-1 (Figure 2). We did not find this correlation in control patients (data shown in supplemental data figure 1)

**Figure 2:**
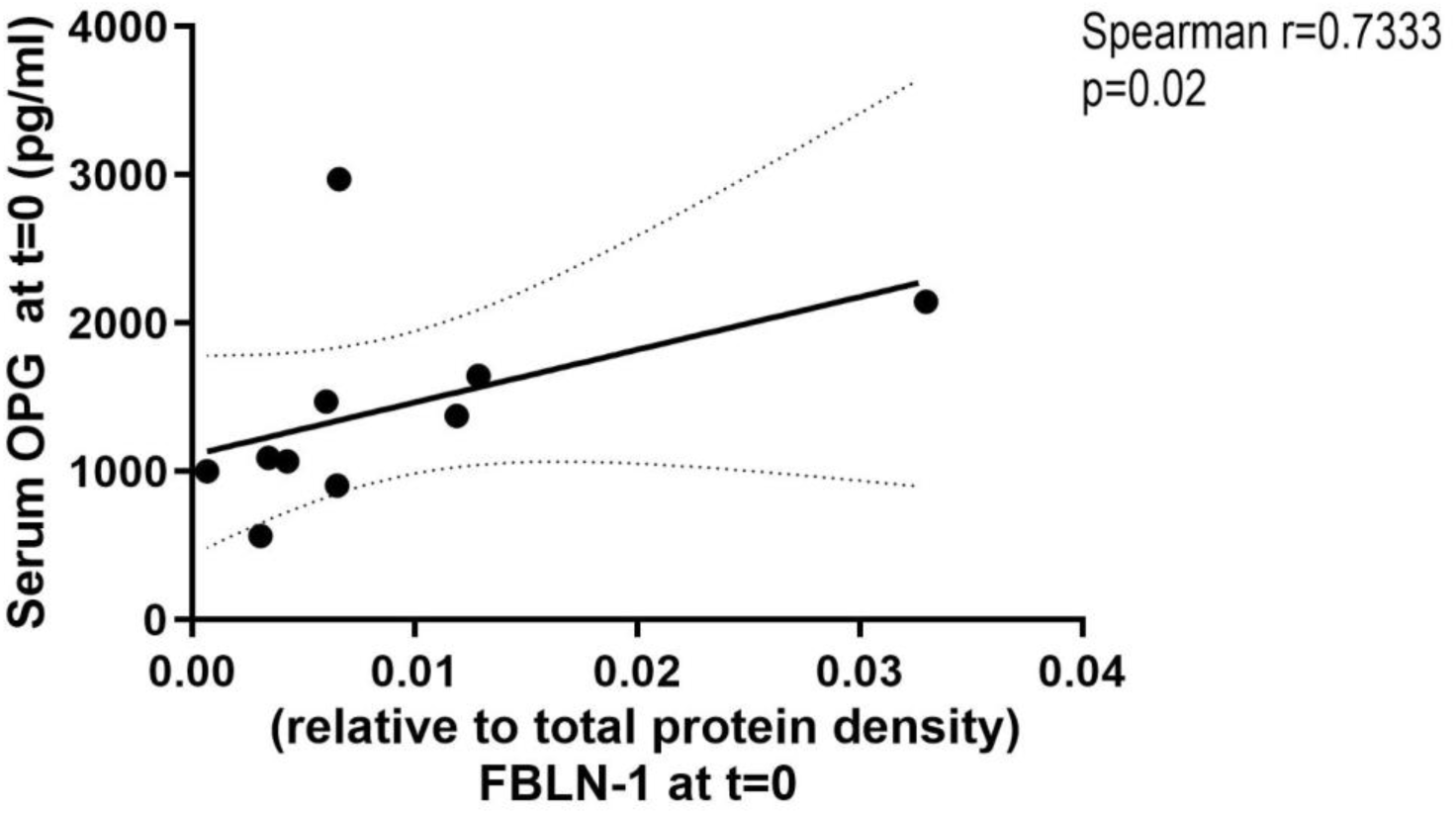
Serum OPG and fibuln-1 levels correlated positively in patients with IPF. Serum of patients with IPF was collected at their first visit to the hospital. Serum from patients with IPF was used to assess OPG using ELISA and fibulin-1 level using western blot^14^. Correlation was tested using a Spearman test (n=10) and p<0.05 was considered significant. OPG: osteoprotegerin; FBLN: fibulin-1

### Tnfrsf11b (OPG) mRNA expression is independent of fibulin-1 in human lung fibroblasts

Since OPG and fibulin-1 were correlating positively in serum from patients with IPF, we continued to investigate whether they may be regulating each other’s expression. To this end, we silenced either Tnfrsf11b (OPG) mRNA or fibulin-1c (Fbln1c) mRNA expression in human lung fibroblasts using siRNA and tested expression of both genes and other fibrosis-associated markers like, collagen-Ⅰ and fibronectin. We found siRNA targeting Tnfrsf11b clearly inhibited Tnfrsf11b expression but it did not influence Fbln1c expression (Figure 3A). Similarly, Fbln1c expression was successfully silenced with siRNA targeting Fbln1c but this silencing did not change Tnfrsf11b expression (Figure 3B). Moreover, mRNA expression of other fibrosis-associated ECM factors were also not changed by downregulating expression of either Tnfrsf11b or fbln1c (supporting Information figure 2).

**Figure 3:**
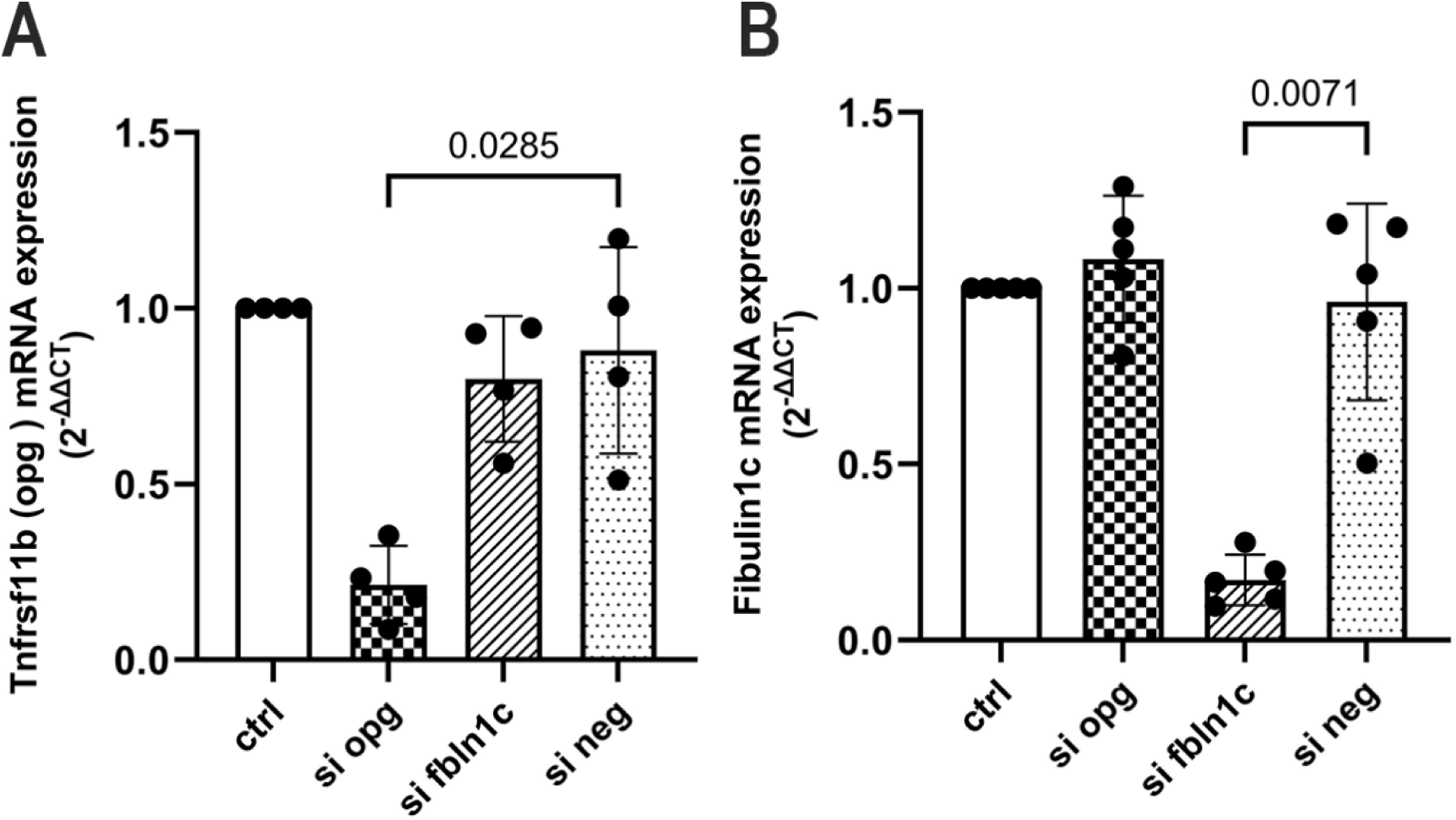
Tnfrsf11b (OPG) mRNA expression is not regulated by fibulin-1 or vice versa in primary lung fibroblasts. Lung fibroblasts were transfected with nothing (ctrl), Tnfrsf11b (OPG) siRNA, fibulin-1c (fbln1) siRNA or scrambled siRNA as a negative control (si neg). Control primary lung fibroblasts cultured for hours, followed by siRNA transfection for 6 hours. After transfection, cells were further to culture for 48 hours. RNA was isolated after total 72 hours culturing. Expression of (A) Tnfrsf11b (n=4) and (B) fibulin-1 c (n=5) was measured by RT-PCR in control lung fibroblasts. Each dot represents an individual patient and data are presented as relative fold gene expression. Data are presented as mean with standard deviation. Groups were compared using a Friedman test and p<0.05 was considered significant. OPG: osteoprotegerin; Fbln1c: fibulin-1c

### OPG protein expression is independent of fibulin-1 in lung fibroblasts

As mRNA does not always reflect protein levels accurately, we further investigated intracellular protein expression of OPG and fibulin-1 in lung fibroblasts of patients with IPF and controls, and in lung fibroblasts isolated from mice with or without the fibulin-1c gene. In human lung fibroblasts, again, silencing of fbln1c inhibited fibulin-1 expression and silencing of Tnfrsf11b clearly inhibited OPG expression and (Figure 4 A-E). However, OPG and fibulin-1 did not influence expression of each other. Additionally, the extracellular secretion of OPG was also found to be independent of fibulin-1c (Figure 4 F, G). A similar pattern was observed for mouse lung fibroblasts, treated with TGFβ1, in which OPG protein expression was not influenced by the absence of fbln1c gene compared to controls with the fbln1c gene (Figure H, I).

**Figure 4:**
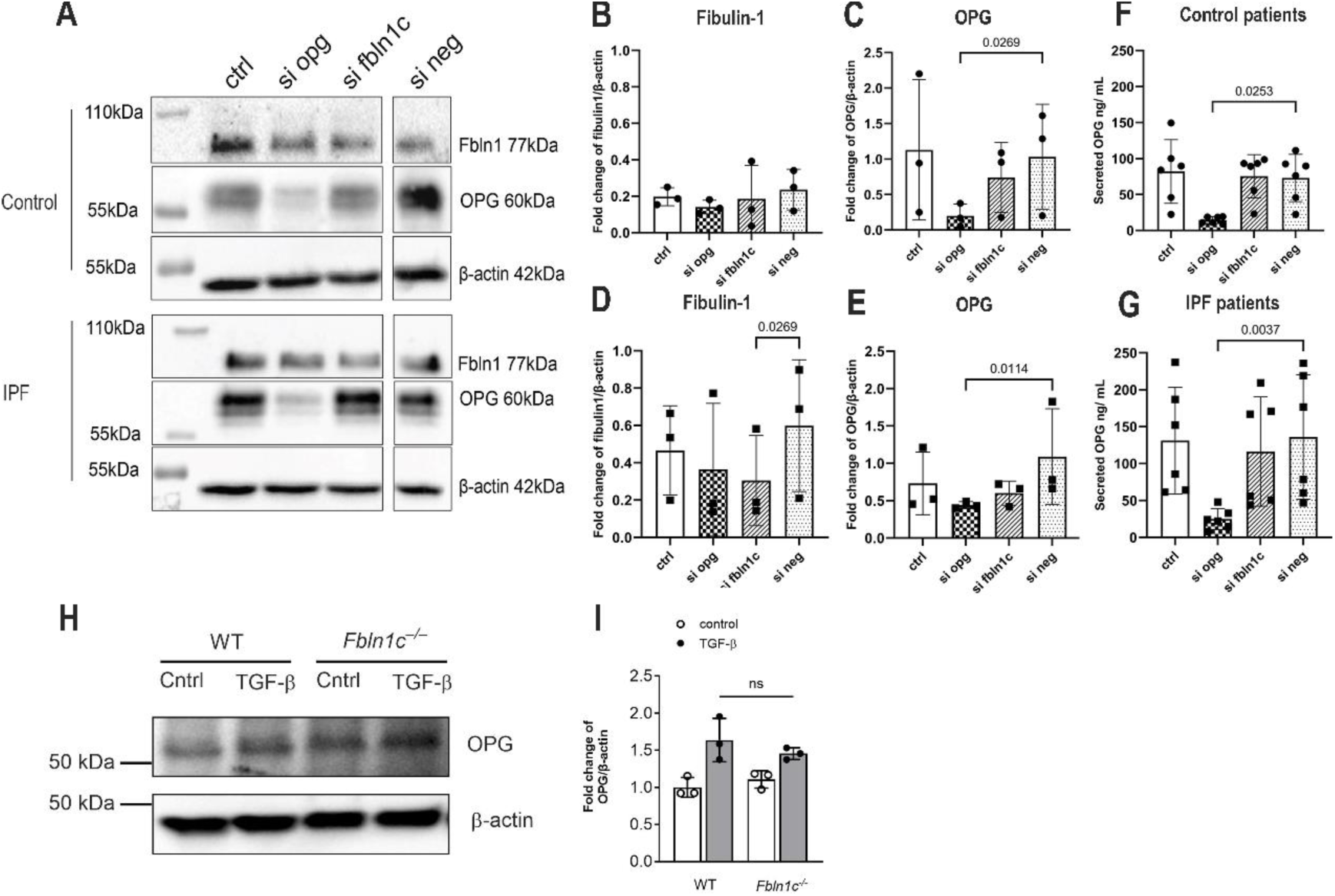
OPG protein expression is neither regulated by fibulin-1 in human lung fibroblasts derived from lung tissue of control patients and patients with IPF nor in mouse lung fibroblasts with or without the fibulin-1c gene and stimulated with TGFβ1. Primary human lung fibroblasts were transfected with nothing (ctrl), Tnfrsf11b (OPG) siRNA, fibulin-1 c siRNA or scrambled siRNA as a negative control. Control primary lung fibroblasts cultured for hours, followed by siRNA transfection for 6 hours. After transfection, cells were further to culture for 48 hours. Cell lysates was isolated after total 72 hours culturing. OPG and fibulin-1 protein levels were measured in cell lysates from control patients (A,) or IPF patients (D) and the quantification from patients shown are shown in (B, C) and from IPF patients are shown in (E, F) (n=3). Protein levels were normalized for Beta-actin levels which was used as loading control, presented as relative fold protein expression. Secreted OPG (G,H) was measured in supernatants from cell cultures of control and fibrotic fibroblasts (n=6). Each dot represents an individual patient. (I, J) OPG expression in lung fibroblasts isolated from mice, with or without the fibulin-1c gene, upon stimulation with TGFβ1 (n=3). Each dot represents an individual mouse and data are presented as relative fold protein expression. Data are presented as mean with SD. For human lung fibroblasts groups were compared using a Friedman test and a Kruskal-Wallis test was used to compare different groups of mouse lung fibroblasts. P<0.05 was considered significant. IPF: idiopathic pulmonary fibrosis; OPG: osteoprotegerin; Fbln1c: fibulin-1c; TGFβ: transforming growth factor beta.

### Less OPG deposition in lung tissue of Fibulin-1 c**^−/−^**mice

Since we found that OPG expression is independent from fibulin-1 in cells, we subsequently questioned if they could influence each other’s deposition in the extracellular microenvironment. To address this question, immunohistochemistry was used to investigate the presence of OPG in lung tissue in fibulin-1c deficient mice and wildtype controls. To induce lung fibrosis both groups of mice were treated with bleomycin, while control animals in each group were treated with saline. Using a two-way ANOVA to analyze the data we found that the effects of bleomycin treatment and presence of fibulin-1c interacted significantly, meaning we could not interpret the outcomes for the separate effects anymore in this test. We therefore did a post-hoc analysis to explain the interaction and we found a trend towards more OPG deposition in lung tissue after bleomycin when compared to saline-treatment in wildtype mice, but this phenomenon was not observed in mice in which fibulin-1 c was deleted (Figure 5 A, B). These data indicate that more OPG deposits in lung tissue of mice with bleomycin-induced lung fibrosis compared to saline-treated controls in wildtype animals, but this deposition is gone in the absence of fibulin-1c.

**Figure 5.**
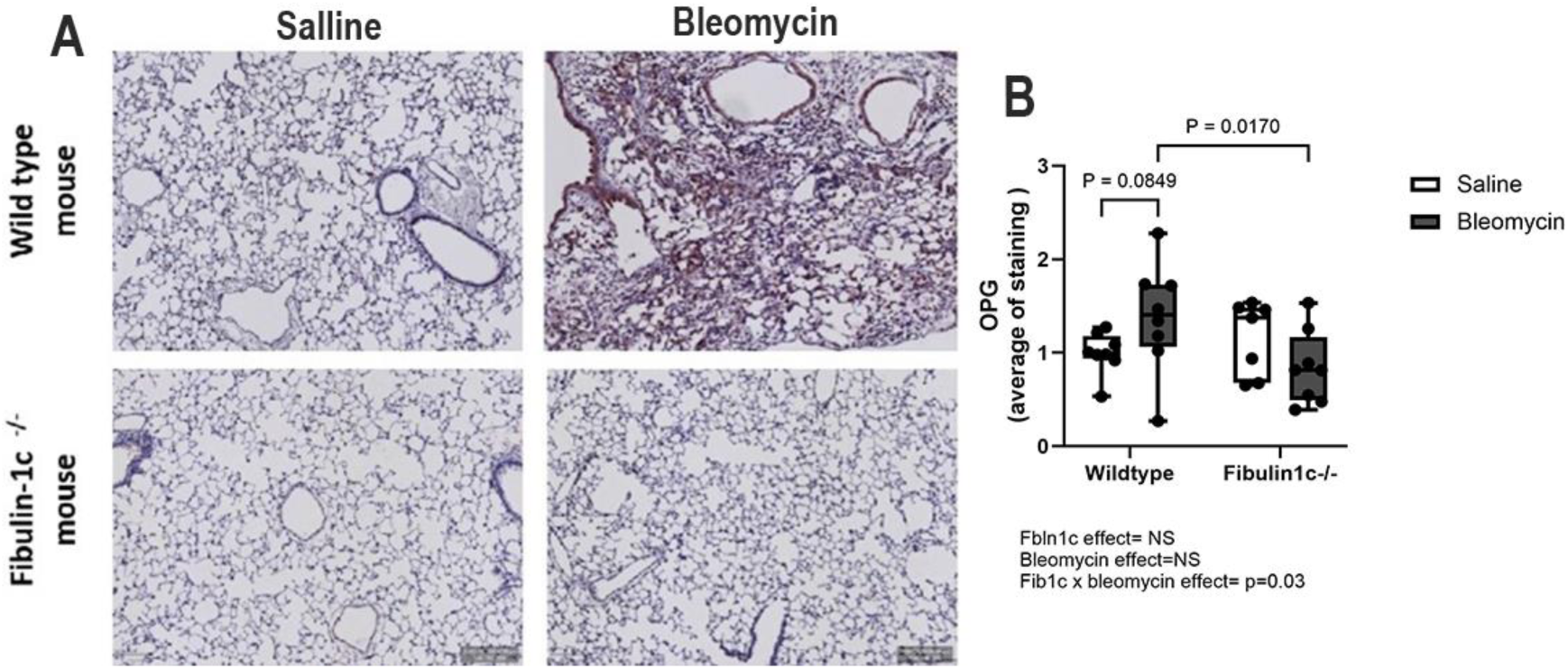
Fibulin-1c deficiency results in less OPG in lung tissue of bleomycin-treated mice. A single administration of bleomycin was used to induce pulmonary fibrosis in wildtype and fibulin 1c-deficient (fibulin-1 c−/−) mice that were sacrificed 28 days later. Control mice received PBS. (A) Representative images of lung tissue from both saline and bleomycin-exposed wildtype and fibulin-1 c−/− mice stained for OPG, scale bar = 300 µm. (B) Quantification of the average of positive OPG staining within the whole tissue (n= 7 or 8 mice). Each dot represents a mouse, and data are presented as the average of percentage of positive OPG-stained within the whole tissue. Groups were compared using a two-way ANOVA with Fisher Least Significant Difference post-hoc test and p<0.05 was considered significant. OPG: osteoprotegerin.

**Figure 6.**
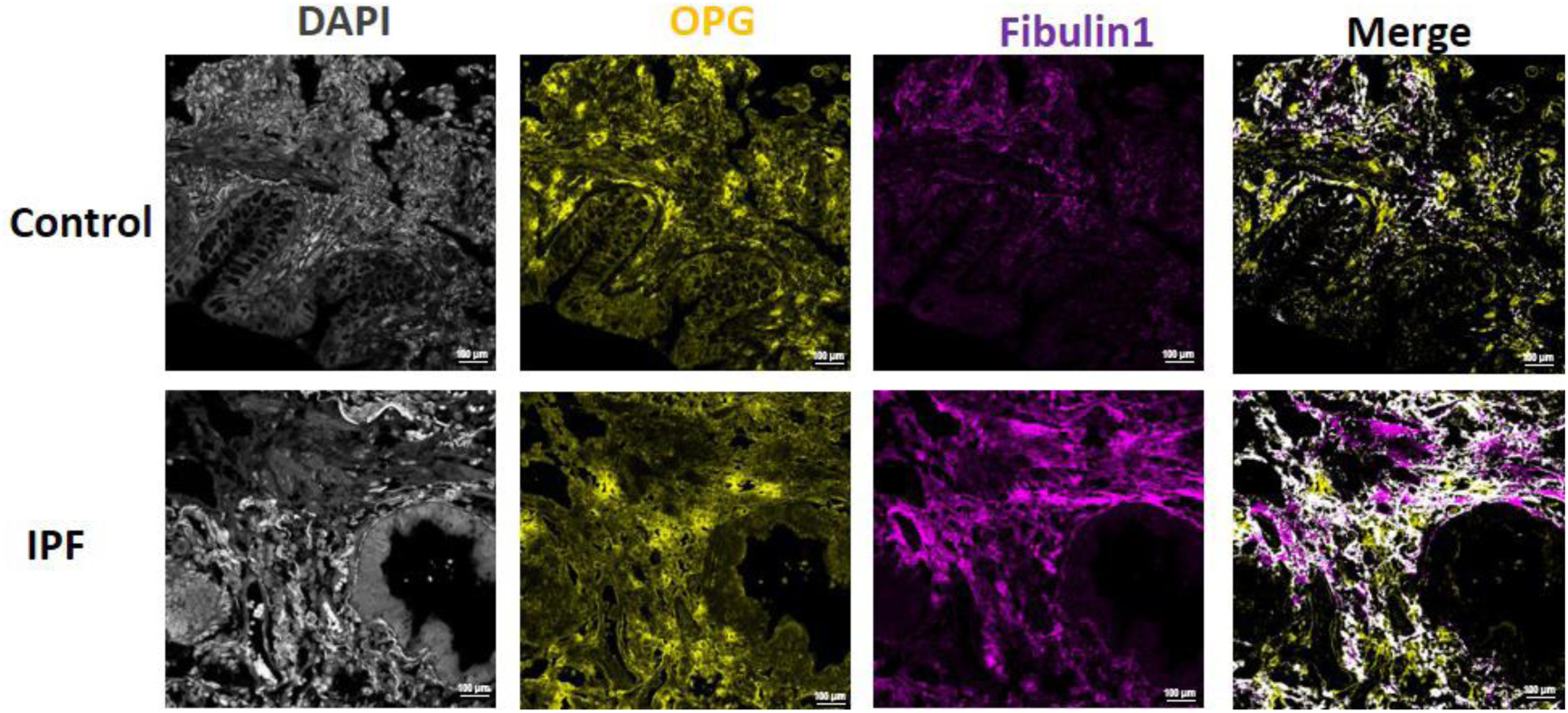
OPG and fibulin-1 distribution in lung tissues from controls and patients with IPF. Immunofluorescent staining was used to identify colocalization of OPG and fibulin-1 in lung tissue from controls and patients with IPF. Representative images show OPG in yellow, fibulin-1 in magenta and colocalization of the two in white, scale bar = 100 µm. Images are representative of staining seen in x control and x IPF patients lung tissues. OPG: osteoprotegerin; IPF: idiopathic pulmonary fibrosis.

### OPG and Fibulin-1 colocalize in lung tissue from control patients and patients with IPF

Given our finding indicated an effect of fibulin-1c deletion on OPG deposition in mouse lung tissue, we decided to investigate their interaction in human lung tissue. To this end, we used immunofluorescence to characterize the deposition of OPG and fibulin-1 separately in lung tissue and also investigated if they colocalize. We observed that OPG was mainly distributed in epithelium and interstitium in both control lung tissue and tissue from patients with IPF, while fibulin-1 was mainly found in the basement membrane and interstitial regions in both types of lung tissue. Colocalization of OPG and fibulin1 was predominantly seen in interstitial regions. These results suggest that OPG and fibulin1 are distributed somewhat differently, but can indeed colocalize together in lung tissue from both controls and patients with IPF.

### OPG, fibulin-1 and LTBP1 colocalize in control and IPF lung tissue

Our previous study reported a potential novel profibrotic mechanism of fibulin-1 in lung fibrosis, showing that it can bind to latent TGF-β-binding protein 1 (LTBP1), regulating activation of TGFβ1 and promoting fibroblasts to increase production of ECM proteins, therefore contributing to fibrogenesis in IPF ^17^. In this study we were therefore curious whether OPG, fibulin1 and LTBP1 may interact in some way. For this purpose, we did immunofluorescent triple staining of OPG, fibulin1 and LTBP1 in control lung tissue and lung tissue from patients with IPF. We observed that LTBP1 is distributed similarly as OPG, i.e. in epithelium and interstitial area. Colocalization of these three proteins was also located in interstitium in both lung tissue from both control and patients with IPF.

### Close proximity of OPG and Fibulin-1, Fibulin-1 and LTBP1 in both control and IPF lung tissue

Colocalization in immunofluorescence does not necessarily mean the proteins are genuinely interacting. Therefore, we performed a proximity ligation assay to determine if any pair of the three proteins would be close enough (<40 nm) for interactions (Figure 7). Here we found that OPG and fibulin-1, as well as fibulin-1 and LTBP1 were located close enough to generate positive staining, particularly in interstitial regions but not in airways or vessels. Of note, we found no close association between OPG and LTBP1. We also noted that the signal of proximity between OPG and fibulin1, as well as fibulin1 and LTBP1 was strongly present in control lung tissue, but more weakly so in fibrotic lung tissue. Combining our immunofluorescence data and proximity ligation data, we show that OPG and fibulin-1 are indeed close enough for physical interactions, as are fibulin-1 and LTBP1, but OPG and LTBP1 are not.

**Figure 7.**
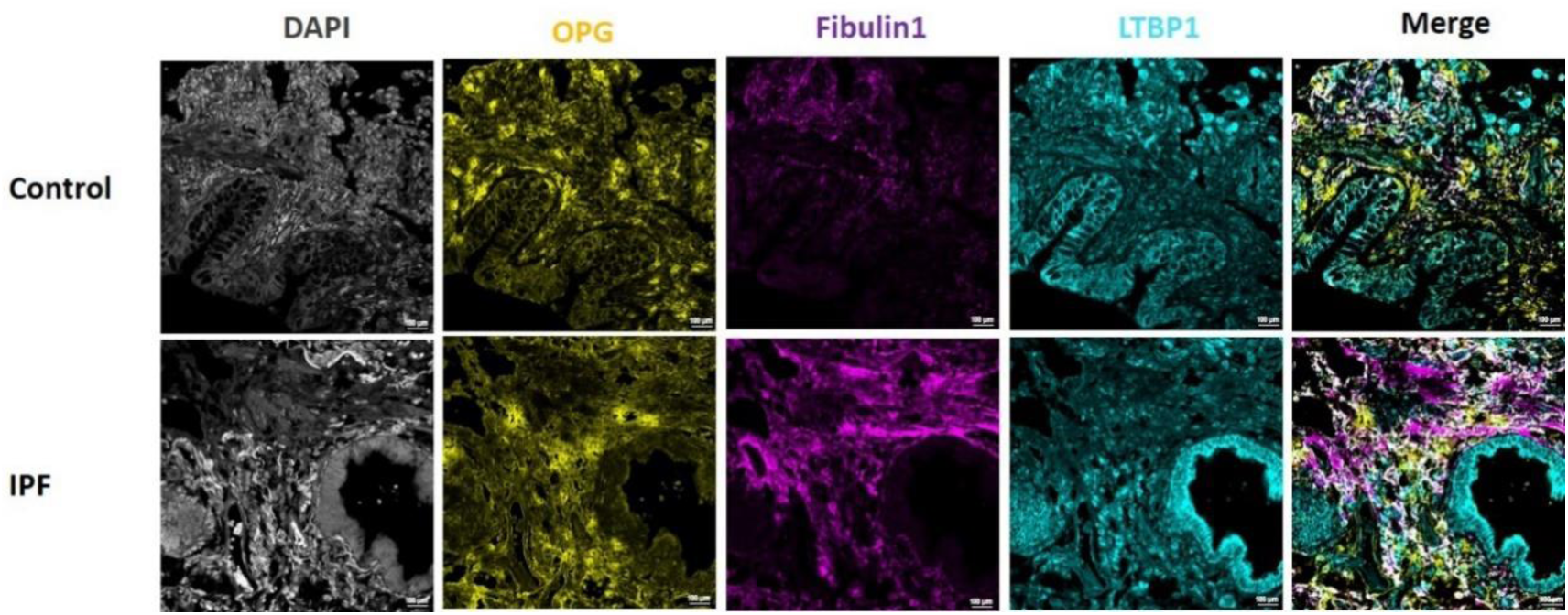
OPG, fibulin-1 and LTBP1 distribution in lung tissue from controls and patients with IPF. Lung tissue sections from controls and patients with IPF were stained using immunofluorescence to identify colocalization of OPG, fibulin-1 and LTBP1. Representative images show the OPG in yellow), fibulin-1 in magenta, LTBP1 in cyan and colocalization in white, scale bar = 100µm. Images are representative of staining seen in x control and x IPF patients lung tissues. IPF: idiopathic pulmonary fibrosis.; OPG: osteoprotegerin; LTBP1: latent TGFβ1 Binding Protein 1.

**Figure 8:**
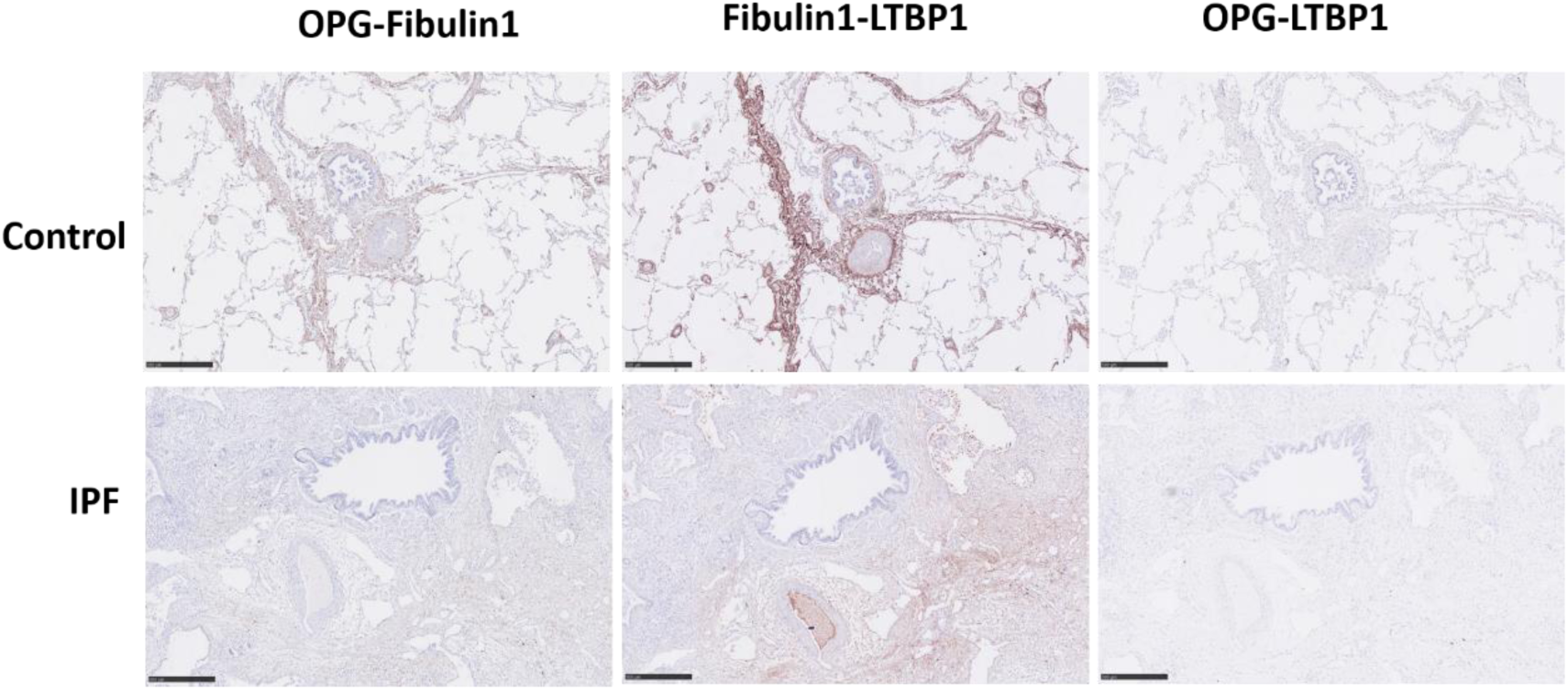
Proximity staining of OPG, fibulin-1 and LTBP1 in lung tissue from both controls and patients with IPF. Representative images of proximity detection of OPG and fbln1, OPG and LTBP1, fbln1 and LTBP1 in lung tissue obtained from control patients and patients with IPF, scare bar = 500 µm. Each two of primary antibodies of OPG, fbln1 and LTBP1 were simultaneously incubated in control lung tissue (n=5) and IPF lung tissue (5). Images are representative of staining seen in x control and x IPF patients lung tissues. IPF: idiopathic pulmonary fibrosis; OPG: osteoprotegerin; LTBP1: latent TGFβ1 Binding Protein 1.

## Discussion

In this study, we initially observed a positive correlation between levels of OPG and fibulin-1 in serum of patients with IPF, suggesting a potential interaction between these proteins in the pathogenesis of IPF. Based on this observation, we hypothesized that OPG may contribute to development of lung fibrosis via an interaction with fibulin-1. However, using gene silencing we convincingly showed that these interactions do not take place intracellularly in fibroblasts as silencing either OPG or fibulin-1 did not change expression of the other protein. Interestingly, in vivo silencing of fibulin-1 did lead to less deposition of OPG in lung tissue from mice exposed to bleomycin compared to controls, suggesting that fibulin-1 is important for retaining OPG in the extracellular milieu. To further explore their potential interaction, we focused on human lung tissues and found that OPG and fibulin-1 extensively colocalized in lung tissue from both control patients and patients with IPF. Since one of our previous studies demonstrated that fibulin1 interacts with LTBP1 we further investigated the relationship between these three proteins, showing that OPG, fibulin-1 and LTBP1 all colocalize in human lung tissues. Proximity ligation assays confirmed a close association between OPG and fibulin-1, as well as fibulin-1 and LTBP1, but not between OPG and LTBP1. Taken together, our data suggest that OPG, fibulin-1 and LTBP1 form a complex in the extracellular space, which may influence development and/or progression of pulmonary fibrosis.

A variety of fibrosis-associated proteins have been reported in fibrotic lung disease {refs}. OPG is among those shown to be associated with different fibrotic diseases^27,28,29^. Our own previous data and those from Bowman et al. suggest that OPG may be a biomarker for progressive lung fibrosis^30^. Nonetheless, the role of OPG in fibrotic lung disease is still unknown. Given the correlation between OPG and fibulin-1 we found in serum of IPF patients and the fact that both proteins are produced by fibroblasts, we focused our attention on these cells. However, our data clearly indicated these proteins do not regulate each other’s expression in fibroblasts. Instead, we found an extracellular interaction, which is not surprising as OPG and fibulin-1 both exhibit a propensity for protein binding. As a secreted decoy receptor of TRAIL and RANKL, OPG is naturally prone to bind these proteins with specificity^31^. However, OPG has also been shown to interact with many other extracellular proteins, including connective tissue growth factor (CTGF), C-telopeptide of type I collagen and glycosaminoglycans^32,33,34^. Fibulin-1 on the other hand is a glycoprotein and is intermixed into the ECM produced by fibroblasts, playing a key role in regulating ECM structure. Numerous studies have found that fibulin-1 interacts with multiple matrix proteins, such as collagen I, fibronectin, laminin, and elastin among others^35,36,37^. In our study, we now show OPG can be added to the list of proteins interacting with fibulin-1 in extracellular space and that OPG deposition in the extracellular matrix relies on the presence of fibulin-1.

Furthermore, we used a combination of immunofluorescence and proximity ligation to further elucidate these interactions, also building on the fact that fibulin-1 interacts with LTBP1. Immunofluorescence shows that OPG, fibulin-1 and LTBP1 colocalized together, but this does not automatically mean the proteins actually interact. Proximity ligation on the other hand can detect if two proteins are located within 40 nm ^24^. Proximity ligation assay confirmed close interactions between OPG and fibulin-1, as well as between fibulin-1 and LTBP1. Notably, OPG and LTBP1 were not found in close association, suggesting a three-way interaction with fibulin-1 in the middle and OPG and LTBP1 on either end. Interestingly, the proximity signal between OPG and fibulin-1 in lung tissue from control patients appeared to be stronger than the signal in fibrotic lung tissue. A possible explanation for this observation may be alterations in ECM protein structure in the fibrotic microenvironment, in particular changes in the organization of the collagen fiber network. Especially type Ⅰ collagen is a critical component of IPF-associated matrix remodeling and its structure may possibly impact the OPG-fibulin-1 binding^38^. Collectively, our findings support the hypothesis that OPG-fibulin-1-LTBP1 form a complex with fibulin-1 acting as a bridge between OPG and LTBP1 and that this interaction is distorted when the ECM becomes fibrotic.

Although the role of OPG in lung fibrosis remains elusive, our study demonstrated that extracellular interactions are important for its retention in the lung. Our study also had some limitations that need to be considered. The limited number of patients used for the different elements of the study may result in a potential bias when extrapolating the findings to a broader range of pulmonary fibrosis patients. Additionally, bleomycin-induced injury does not fully mimic human lung fibrosis, which may lead to limited translatability. Nevertheless, we provided a consistent observation that OPG and fibulin-1 interact within the fibrotic extracellular microenvironment in both mouse and human lung tissue and form a complex that may contribute to development and/or progression of lung fibrosis.

In conclusion, our study has identified a novel protein complex consisting of OPG, fibulin-1 and LTBP1, which may play a role in lung fibrosis. Through investigating the (post)translational regulation, distribution, and spatial distance, we show this complex to be present within both control and fibrotic extracellular microenvironments. An interesting next step would be to interfere with this complex to investigate how it contributes to development and/or progression of lung fibrosis.

