## Supplemental Figure 1, Supplemental Figure2 for "Interaction of osteoprotegerin with fibulin-1 is essential for extracellular matrix deposition in fibrotic lung tissue"

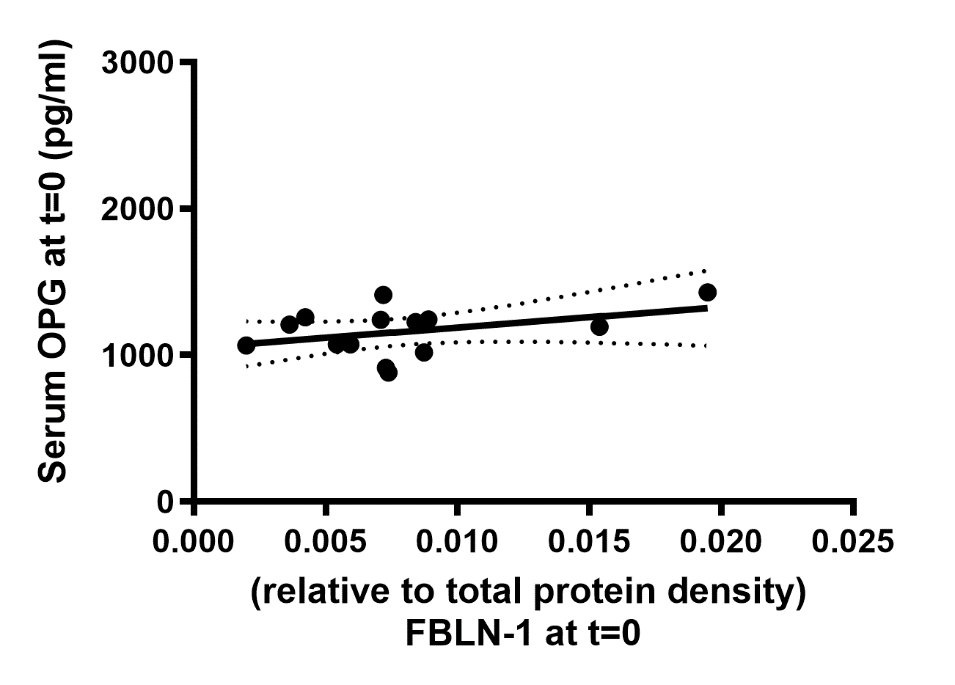

**Figure 1: Serum OPG and Fibuln-1 levels correlated positively in Sserum from non-IPF patients.** Serum of patients without IPF was collected at their first visit to the hospital. Serum from non-IPF patients was used to assess OPG level using ELISA and fibulin-1 level using western blot. Correlation was tested using a Spearman test (n=10) and p<0.05 was considered significant.

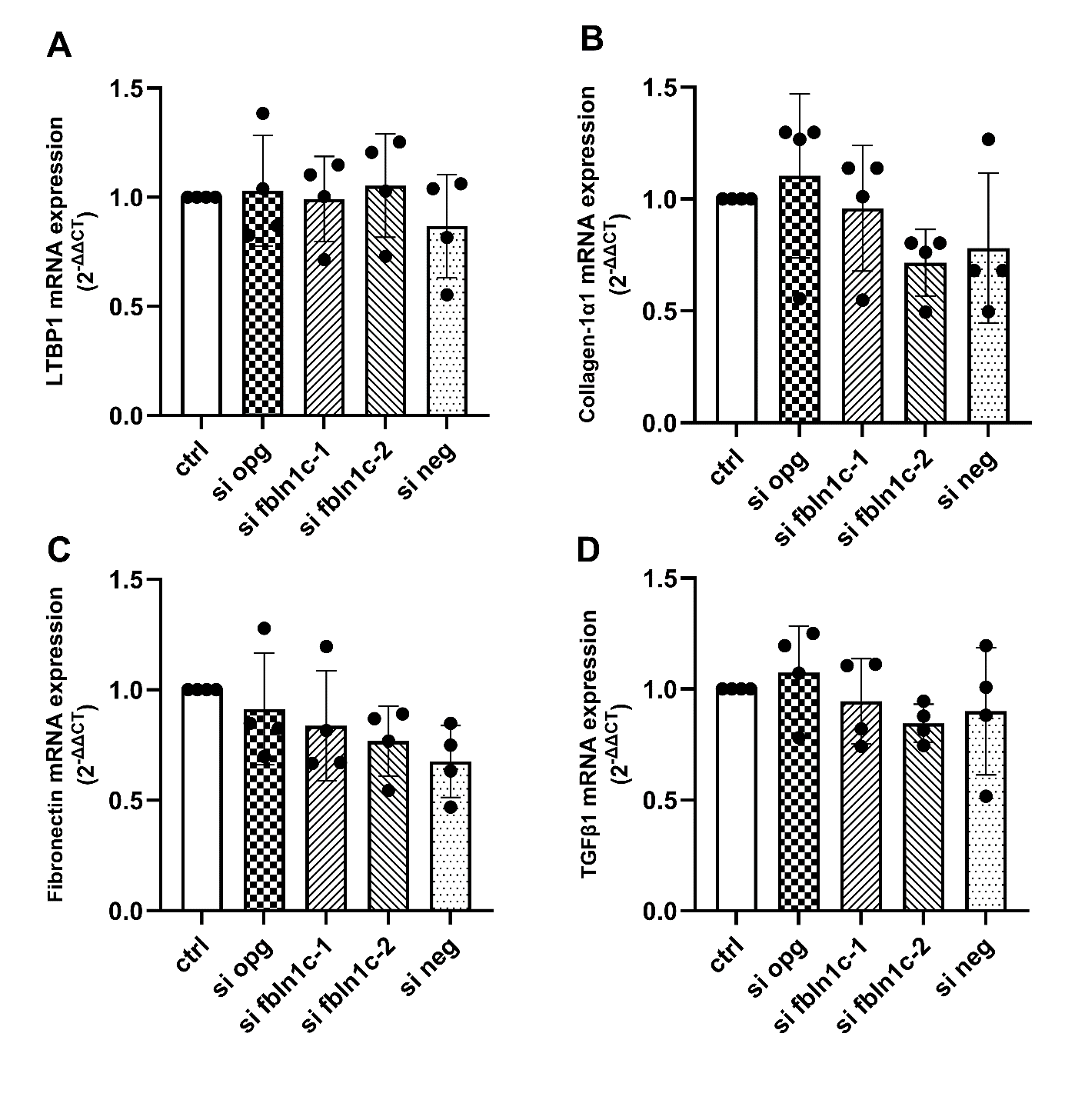
**Figure 2: gene expression of fibrosis-associated ECM factors is not regulated by fibulin-1 and OPG in primary lung fibroblasts.** Lung fibroblasts were transfected with nothing (ctrl), Tnfrsf11b (OPG) siRNA, fibulin-1c (fbln1) siRNA or scrambled siRNA as a negative control (si neg). Control primary lung fibroblasts cultured for hours, followed by siRNA transfection for 6 hours. After transfection, cells were further to culture for 48 hours. RNA was isolated after total 72 hours culturing. Expression of (A) LTBP1 (n=4), (B) collagen 1α1 (n=4), (C) fibronectin (n=4) and (D) TGFβ1 (n=4) was measured by RT-PCR in control lung fibroblasts. Each dot represents an individual patient and data are presented as relative fold gene expression. Data are presented as mean with Standard Deviation. Groups were compared using a Friedman test and p<0.05 was considered significant. ECM: extracellular matrix; LTBP1: latent TGFβ1 Binding Protein 1; TGFβ1: transforming growth factor beta 1.
